# Colorimetric Hydrogel Dressing with Smartphone Detector for Point-of-Care Wound pH Monitoring

**DOI:** 10.64898/2026.08.07.742070

**Authors:** Katia Cherifi, Katerina Christodoulopoulos, Aylin Kizilkaya, Mariam Touba Toure, Farnoush Toupchinejad, Simon Matoori

## Abstract

Chronic wounds such as diabetic foot ulcers are typically more alkaline than healing wounds, making wound pH a valuable diagnostic and prognostic marker. However, point-of-care pH monitoring remains limited by the availability of point-of-care wound pH sensing systems that offer quantitative pH determination, low toxicity, and small portable detectors. Here, we report a colorimetric pH-sensing wound dressing that enables *in situ* pH detection using a conventional smartphone camera. The anionic pH-sensitive dye HPTS was loaded onto cationic microparticles and embedded within a calcium-crosslinked alginate hydrogel. Across the clinically relevant range of pH 6.0–9.0, increasing pH produced a progressively more intense yellow coloration, quantified through the blue channel of smartphone-acquired RGB images. The dressing displayed a strong, rapid, and reversible signal *in vitro* with low dye release. In a full-thickness excisional wound model in mice, wound pH changes were detected *in vivo*. The combination of a pH-sensitive colorimetric hydrogel with a conventional smartphone detector offers accessible wound pH monitoring at the point-of-care.

## 1. Introduction

Diabetic foot ulcers (DFUs) are among the most common chronic lower-extremity wounds and constitute a serious complication of diabetes, affecting up to 34% of individuals with diabetes. ^1, 2^ These wounds heal slowly, with only approximately half closing completely within seven months, and they recur frequently, in up to 60% of cases within three years. ^2^ DFUs are the leading cause of lower-limb amputation worldwide and are associated with amputation in approximately 20% of affected patients. ^1, 3^ Despite an expanding understanding of the molecular pathophysiology of diabetic wounds, current clinical guidelines still rely on macroscopic assessment based on ulcer depth, involvement of underlying tissues, and non-specific inflammatory indicators such as redness and swelling. ^4–7^ Given the absence of molecular diagnostics for DFU staging and prognosis, there is strong interest in point-of-care technologies for chronic wound diagnosis and monitoring. ^8–10^ As molecular therapies for DFUs emerge, such diagnostics may further enable selection of the most appropriate treatment for individual patients and evaluation of therapeutic success.

Wound pH has been identified as a relevant therapeutic target in chronic wounds. ^1, 11–16^ Chronic wounds, including diabetic ulcers, tend to exhibit a more alkaline pH (7.5–9.0) than normally healing wounds, which are slightly acidic and approach the pH of healthy skin (near pH 6.0). ^1, 15–18^ This elevated pH reduces wound oxygenation, reduces wound oxygenation, increases proteolytic enzyme activity, and favors the growth of pathogenic bacteria, ^19^ and promotes macrophage polarization toward the pro-inflammatory (M1) phenotype. In contrast, an acidic-to-neutral pH attenuates chronic inflammation and bacterial growth and promotes polarization toward the pro-healing (M2) phenotype. ^19–21^ A low wound pH further establishes favorable conditions for pro-healing processes, including angiogenesis and the migration and proliferation of keratinocytes and fibroblasts. ^22, 23^ Because pH is distributed heterogeneously across the wound bed, it is an attractive target for real-time, spatially resolved monitoring of wound healing.^17, 24^

We previously reported a fluorescent pH-sensing bandage for point-of-care detection of wound pH. ^25^ The system employs a double-encapsulation strategy to minimize dye leakage into wound tissue: a pH-sensitive fluorescent dye is loaded onto microparticles, which are in turn immobilized within a hydrogel matrix. The anionic pH-sensitive dye HPTS is adsorbed onto cationic microparticles, and the dye-loaded microparticles are encapsulated within ionically crosslinked calcium-alginate hydrogels. As wound fluid enters the hydrogel, the fluorescence profile of the dye changes as a function of pH. In a clinical setting, this fluorescence signal would be quantified with a portable fluorometer for point-of-care pH sensing. However, because portable fluorometers remain an emerging technology with limited availability and relatively high cost, diagnostic systems based on more accessible detection methods such as colorimetry are needed. ^26, 27^

Colorimetric sensors based on pH indicators are simple, reliable, and enable rapid pH measurement owing to fast proton-exchange kinetics. ^26–31^ In this work, we exploited the colorimetric properties of HPTS, the pH-sensing dye used in our fluorescent pH-sensing bandage, at increased concentrations to enable pH quantification by RGB colorimetry (Figure 1). The RGB color model is an additive color system used in digital displays and imaging, in which colors are produced by combining different intensities of red, green, and blue light. ^27, 32^ The absorbance of HPTS shifts from colorless/pale yellow at acidic pH to bright yellow at alkaline pH, with a pKa of approximately 7.5. ^33–38^ Building on this pH-dependent absorbance behavior, we developed a second-generation, colorimetric pH-sensing bandage and evaluated it using a smartphone camera to enable point-of-care detection without the need for a dedicated detector.

**Figure 1.**
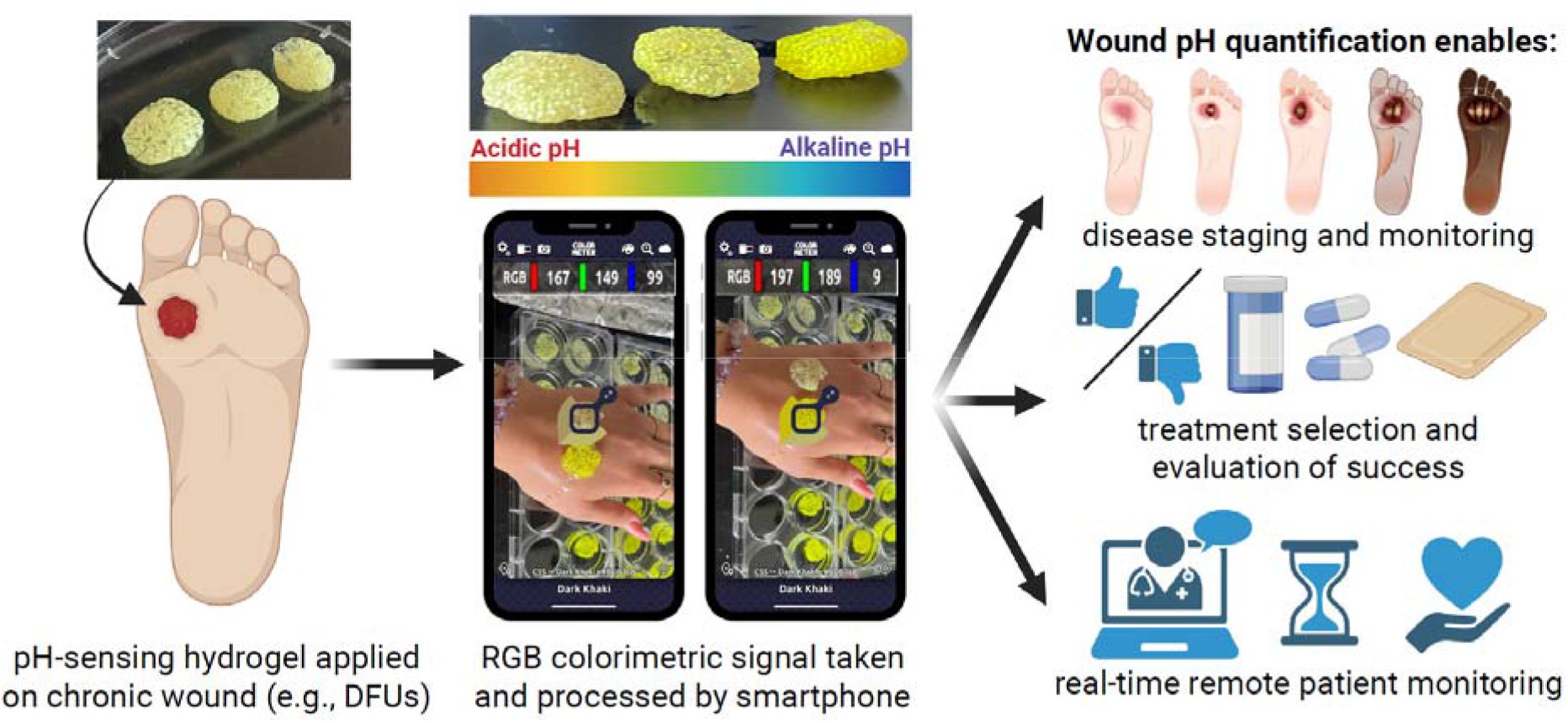
Overview of the colorimetric pH-sensing wound dressing. The hydrogel dressing is applied onto a chronic wound where it absorbs wound exudate and changes its color in response to wound fluid pH. This colorimetric shift is captured with a conventional smartphone camera and quantified via RGB analysis, translating wound pH into an actionable, quantitative readout that enables disease staging, treatment selection, evaluation of treatment success, and remote patient monitoring, with the potential to improve patient outcomes and quality of life.

## 2. Experimental Section

### Materials

8-Hydroxypyrene-1,3,6-trisulfonic acid trisodium salt (HPTS, also known as pyranine), sodium chloride, tris(hydroxymethyl)aminomethane (TRIS) base, 2-(N-morpholino)ethanesulfonic acid (MES), potassium dihydrogen phosphate, Amberlite IRA-900 microparticles (styrene–divinylbenzene matrix with benzyltrialkylammonium [BTA] functionality; BTA MPs), medium-viscosity sodium alginate, calcium sulfate dihydrate, and Whatman Filter papers were all purchased from Sigma-Aldrich (St. Louis, MO, USA). Calcium chloride anhydrous was purchased from Thermo Scientific Chemicals (Waltham, MA, USA).

Disposable biopsy punch tools (Integra Miltex, 6 and 10 mm diameter) and glass microscope slides (UltidentBrand, 75 × 25 × 1 mm) were purchased from UltiDent Scientific (Montreal, QC, Canada). The ColorMeter RGB Colorimeter application (White Marten GmbH) was obtained from the Apple App Store and used on an iPhone 13 (Apple Inc.) as the primary RGB detection platform.

### Methods

#### 2.2.1 Smartphone-Based RGB Imaging Setup

A smartphone-based RGB imaging setup was used to quantify the colorimetric response of HPTS in solution, after loading onto microparticles, and within hydrogels. All measurements were performed with an iPhone 13 (Apple Inc., USA) using the rear camera at a fixed 2× zoom. For reproducible acquisition, the smartphone was mounted on a rigid holder perpendicular to the sample plane at a fixed distance of 30 cm from the plate surface (**Supporting Figure 1A**). Measurements were performed in a darkened environment, with the plate and camera enclosed in a black cardboard box to minimize ambient light and specular reflections. Images were captured with the default iPhone Camera application and imported into the ColorMeter RGB Colorimeter App (White Marten GmbH), ^39^ which extracted mean red (R), green (G), and blue (B) channel intensities from a region of interest (ROI) centered on each sample **(Supporting Figure 1B).** Unless otherwise specified, all measurements were performed in triplicate.

#### 2.2.2 Colorimetric Analysis of HPTS Solutions

To evaluate the concentration-dependent response of the free dye, HPTS was dissolved in isotonic phosphate-buffered saline (PBS, 50 mM, pH 7.4) at 0, 10, 50, 100, 250, and 500 µM. Aliquots were transferred to flat white-bottom 96-well plates (Corning, 353296) and imaged as described in Section 2.2, with the ROI adjusted to cover the entire well area (**Supporting Figure 2A**).

To characterize the pH-dependent response of the free dye, HPTS was prepared at 100 µM in isotonic buffers adjusted to pH 6.0–9.0. MES isotonic buffer (50 mM) was used for pH 6.0 and 6.5, and isotonic TRIS-buffered saline (TBS, 50 mM) for pH 7.0, 7.5, 8.0, 8.5, and 9.0, with pH adjusted using HCl or NaOH as required. Solutions were dispensed into flat white-bottom 96-well plates and analyzed under the same imaging conditions, with RGB intensities extracted from an ROI matched to the well diameter (**Supporting Figure 2B**).

#### 2.2.3 Preparation and Colorimetric Response of HPTS-Loaded Microparticles

In contrast to our previous fluorescent pH-sensing hydrogel, ^25^ we increased the BTA MP content from 5 to 150 mg per alginate hydrogel improved microparticle distribution and enabled a robust signal across the entire bandage surface (**Supporting Figure 3A**). A loading of 150 mg of BTA MPs was selected for *in vitro* testing, corresponding to a hydrogel disk with an area comparable to that of a typical human chronic ulcer (approximately 2–3 cm in diameter). ^1, 40^

For smartphone-based analysis of HPTS-loaded MPs in suspension, 150 mg of BTA MPs was incubated in 5 mL of HPTS solution (0, 100, or 500 µM in PBS, 50 mM, pH 7.4) for 15 min at 37°C under agitation and protected from light. Suspensions were purified from unbound dye by centrifugation (three times at 5,000 × *g* for 1 min) and resuspended in the corresponding buffer. Suspensions were then transferred into 24-well plates in buffers at pH 6.0 to 9.0 (Sarstedt, 83.3921, flat bottom, transparent), allowed to stabilize for 5 min, and imaged as described in Section 2.2 (**Supporting Figure 3B**). Mean R, G, and B values were extracted from each well.

#### 2.2.4. Preparation of HPTS-MP-Loaded Alginate Hydrogels

To load the dye, 150 mg of BTA MPs was incubated in 5 mL of 100 µM HPTS solution (PBS, 50 mM, pH 7.4) for 15 min at 37 °C under agitation and protected from light. Unbound dye was removed by centrifugation (three times at 15,000 × *g* for 1 min); after the final wash, the supernatant was discarded, and the pellet of HPTS-loaded MPs was used for hydrogel preparation.

The pellet was mixed with 240 µL of medium-viscosity sodium alginate solution (8% w/v in water) and 80 µL of calcium sulfate dihydrate suspension (174 mM) according to a previously established protocol, ^25^ yielding final concentrations of 6% (w/v) alginate and 43.5 mM calcium sulfate. The mixture was shaped into disks of approximately 1 x 1 cm with spatulas and transferred into 25 mM CaCl_₂_ solution (100 mL) for 4 h at 37 °C for ionic crosslinking. The resulting hydrogels contained 150 mg of HPTS-loaded MPs homogeneously distributed throughout the alginate matrix (**Supporting Figure 4A**).

#### 2.2.5 *In Vitro* and *Ex Vivo* pH-Sensing

For *in vitro* measurements, hydrogel disks were placed in 6-well plates (Sarstedt, 83.3920) and incubated in 3 mL of isotonic buffers adjusted to pH 6.0–9.0 (MES 50 mM for pH 6.0–6.5; TBS 50 mM for pH 7.0–9.0) RGB images were recorded at 0, 2, 3, 5, 10, 15, 30, 60, and 120 min after immersion, with mean R, G, and B values extracted from an ROI encompassing the entire hydrogel surface. The B channel intensity served as the primary kinetic readout. Signal stability was assessed by monitoring the B channel over 120 min and determining the time to plateau. Reversibility was evaluated by sequentially transferring the same hydrogels between pH 6.0 and pH 9.0 buffers, changing the buffer every 4 min while recording the B channel continuously. All kinetic experiments were performed at room temperature in triplicate.

Hydrogels were equilibrated for 5 min at room temperature to allow proton diffusion and signal stabilization, then imaged as described in Section 2.2. Mean R, G, and B values were extracted from an ROI covering the entire visible hydrogel area; the B channel intensity served as the primary readout of pH.

For *ex vivo* measurements, full-thickness excisional dorsal wounds were created in mice using a 10 mm biopsy punch (C57BL/6 mice (males, 18 weeks of age) and Podocin-Adam17 mice (albino female, 11 weeks of age), obtained from Charles River (Montreal, QC, Canada), n = 3), and the excised skin was used immediately as an *ex vivo* wound model. Hydrogels pre-equilibrated in 3 mL of buffers of defined pH (6.0–9.0) were placed onto the wound surfaces with full coverage (**Supporting Figure 4A, B**). After 5 min of contact, images were acquired and analyzed as above (**Supporting Figure 4C**). All *in vitro* and *ex vivo* experiments were performed in triplicates Dye release from the hydrogel loaded with HPTS-containing microparticles was assessed in 3 mL of the same buffers over an extended period of 7 days. At each time point, 100 µL of the surrounding buffer was collected and HPTS release was quantified by measuring fluorescence intensity at 413 nm, the isosbestic wavelength of HPTS at which fluorescence is independent of pH, allowing direct quantification of released dye regardless of buffer pH.

#### 2.2.6. *In Vivo* pH Sensing in a Murine Wound Model

Hydrogel Patch Preparation. For the *in vivo* application, the hydrogel was adapted to obtain thin, flat patches with increased microparticle loading and improved conformity to the wound bed, while preserving the same HPTS-loaded BTA MP chemistry. Briefly, 340 mg of BTA MPs was loaded with HPTS in 10 mL of HPTS solution (100 µM in PBS, 50 mM, pH 7.4), purified as above, and freed of residual supernatant. The purified MPs were mixed with 240 µL of medium-viscosity sodium alginate solution (8% w/v) using a spatula, after which 80 µL of calcium sulfate suspension (400 mM) was added and manually mixed. To form thin films, two microscope slides were used as described in the literature; ^41^ the slides were dipped in 100 mM CaCl_₂_ and the mixture placed between them and flattened under gentle compression (225 mL of water in a 250 mL beaker for reproducibility) while submerged in the same CaCl_₂_ solution. Crosslinking proceeded for 10–15 min under this applied weight to yield films approximately 1 mm thick. Circular patches matching the wound dimensions were prepared using a sterile 6 mm biopsy punch; one film yielded approximately 10 wound-sized patches (**Supporting Figure 5A**). To validate the pH sensitivity of this adapted geometry, patches were incubated for 10 min in isotonic MES buffer (50 mM, pH 6.0–6.5) or TBS buffer (50 mM, pH 7.0–9.0) (**Supporting Figure 5B**), and blue channel intensity was measured as described above.

Surgical Procedure. *In vivo* pH sensing was performed in healthy male C57BL/6 mice using a full-thickness dorsal excisional wound model under an approved institutional animal care protocol by Université de Montréal. Mice were 14–28 weeks old at experimentation, with most animals 14–16 weeks old, and obtained from Charles River (Montreal, QC, Canada). On day 1, the dorsal hair was shaved and depilated. On the surgery day, anesthesia was induced and maintained with isoflurane (2.0% induction; 1.0–1.5% maintenance), and the dorsal skin was disinfected by sequential swabbing with alcohol, povidone-iodine, and alcohol. Two full-thickness excisional wounds were created on the dorsum of each mouse with a sterile 6 mm biopsy punch, extending through the skin to the *panniculus carnosus*.

Wound pH Measurement. The left wound was used for wound fluid collection and the right wound for direct application of the pH-sensing hydrogel (**Supporting Figure 6A**). For fluid sampling, based on the literature, a 6 mm Whatman filter paper disk was gently applied to the left wound bed at 0, 1, 3, and 24 h post-wounding to collect exudate for reference pH analysis. ^42–44^ For *in vivo* sensing, an *in vivo*-adapted patch was placed directly onto the right wound at the same time points, ensuring full coverage and intimate contact with exudate for 10 min (5 min on each side, flipped halfway through). Between time points, wounds were covered with a Tegaderm dressing to maintain a moist environment. After wound contact, hydrogels were imaged under controlled lighting using the setup above (**Supporting Figure 6B**). Mean R, G, and B intensities were extracted from an ROI encompassing the visible hydrogel surface, and the B channel intensity was used as the primary readout of local wound pH.

## 3. Results & Discussion

### 3.1 Colorimetric analysis of HPTS in solution

In the first generation of the assay, we identified HPTS as a suitable dye for wound pH sensing due to its fluorescence response in the clinically relevant range of pH 6.0–9.0. ^25, 45^ To assess whether HPTS could be quantified by smartphone-based RGB analysis, its absorbance was first characterized over the clinically relevant range of pH 6.0–9.0. HPTS exhibited a strong absorbance band at 455 nm, whose intensity increased with pH (**Figure 2A**). This band overlaps with the blue channel detected by RGB-based smartphone cameras, supporting the use of RGB analysis for colorimetric sensing (**Figure 2A**). Using the smartphone–ColorMeter setup and at a fixed concentration of 100 µM across pH 6.0–9.0, the solutions became progressively more intensely yellow with increasing pH, reflected by a decrease in B channel intensity (**Figure 2B** and **Supporting Figure 2B**). This decrease in B channel intensity with increasing absorbance in the yellow spectrum is in accordance with the literature. ^31, 32, 46–48^ Together, these results confirm that HPTS provides a robust colorimetric response to pH within the range clinically relevant to chronic wounds, and that this signal can be detected by a smartphone.

**Figure 2.**
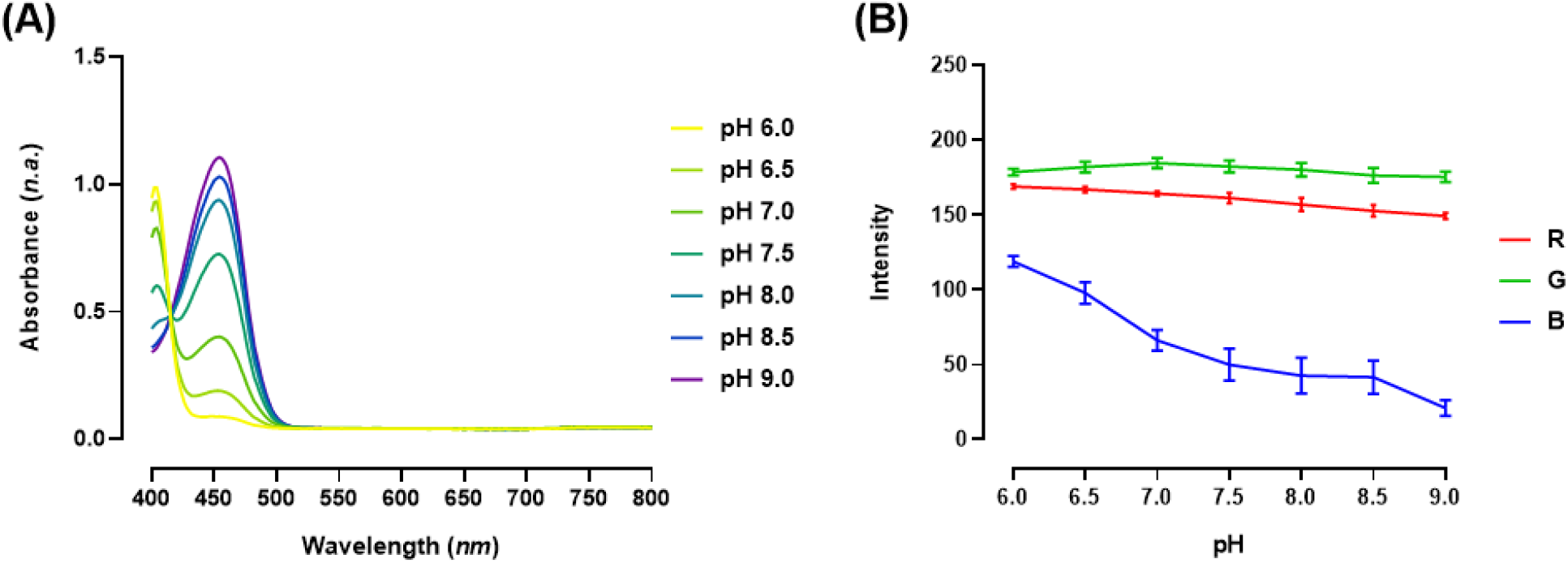
pH-response of HPTS in solution **(A)** Absorbance spectra of HPTS in isotonic PBS buffer (50 mM) at pH 6.0–9.0. **(B)** Red (R), green (G), and blue (B) channel intensities of HPTS solutions (100 µM) at pH 6.0–9.0, extracted from smartphone images using the ColorMeter app. Samples were prepared in MES isotonic buffer (50 mM) for pH 6.0–6.5 and TBS isotonic buffer (50 mM) for pH 7.0–9.0. Data in (B) are presented as mean ± SD (n = 3).

### 3.2 Colorimetric analysis of HPTS-loaded microparticles

To minimize the release of HPTS into the wound tissue, the dye was adsorbed onto microparticles (MPs). In our previous study, we selected cationic BTA microparticles due to the strong binding of the negatively charged HPTS and the preserved pH sensitivity of the dye. ^25^ Adsorbing HPTS onto the MPs anchors the dye through electrostatic interaction between the anionic dye and the cationic particles, which limits its release into the wound once the MPs are immobilized within the final hydrogel dressing. This configuration preserves the sensing chemistry while enabling immobilization within the dressing and minimizing direct exposure of free dye to the wound environment.

Relative to the previously reported fluorescence-based system, ^25^ higher dye loadings were required to generate a colorimetric signal sufficient for absorbance-based smartphone detection. At pH 7.0, increasing the HPTS loading concentration on BTA MPs from 0 to 500 µM produced a clear decrease in the B value, confirming that the HPTS–MP complex could be detected optically with the smartphone–ColorMeter app setup (**Figure 3A**). When these HPTS-loaded MPs were incubated across pH 6.0–9.0, a visible color change was accompanied by a progressive decrease in the B value with increasing pH (**Figure 3B** and **Supporting Figure 3B**).

**Figure 3.**
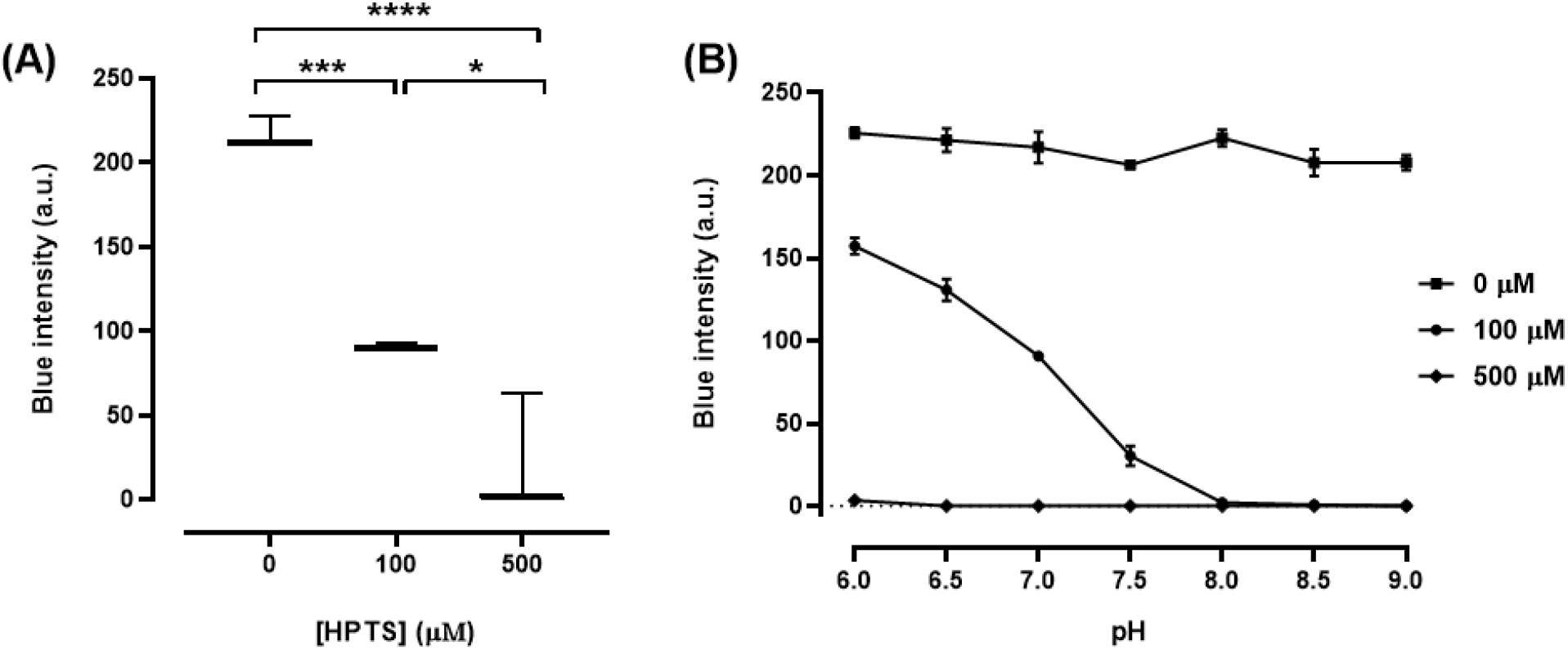
Colorimetric response of HPTS-loaded microparticles. **(A)** Concentration dependence of the blue channel (B value) in HPTS-loaded BTA microparticles at pH 7.0, measured by smartphone-based RGB imaging. **(B)** Blue channel intensity of HPTS-loaded microparticles at different pH values (6.0–9.0). Results are presented as mean ± SD (n = 3). Statistical significance was determined by one-way ANOVA followed by Tukey’s post-hoc test (*p < 0.05, ***p < 0.001, (****p < 0.0001).

Together, these data confirm that the pH-dependent optical response of free HPTS is preserved after adsorption onto BTA MPs. Although the colorimetric format required higher dye loadings than the fluorescence-based system, the B-value signal approached saturation at 500 µM, whereas 100 µM provided a robust, quantifiable response while preserving dynamic range across pH 6.0–9.0 (**Figure 3B**). A loading concentration of 100 µM HPTS, combined with B-value analysis, was therefore selected as the optimized sensing condition for subsequent hydrogel preparation and testing.

### 3.3. *In vitro* activity and stability of HPTS MPs-loaded alginate hydrogel

To obtain a wound-compatible colorimetric dressing, HPTS-loaded BTA MPs were encapsulated in a calcium-crosslinked alginate hydrogel. This hydrogel was chosen for its nanoporous architecture, which effectively confines the microparticles within the hydrogel scaffold. Calcium alginate was also selected because of its high biocompatibility and low immunogenicity. ^49–51^ Notably, calcium-crosslinked alginate hydrogels have received regulatory approval for use as wound dressings in DFUs. ^52–55^ Because smartphone-based RGB analysis averages the signal over the visible surface of the sample, a homogeneous distribution of microparticles throughout the hydrogel was essential for robust readout. Increasing the microparticle content from 5 mg, the loading used in the previously reported fluorescent bandage, ^25^ to 150 mg per hydrogel improved spatial distribution within the matrix and enabled signal detection across the full hydrogel surface (**Supporting Figure 3A**). Hydrogels incubated in buffers spanning pH 6.0–9.0 exhibited a clear, time-dependent decrease in B channel intensity with increasing pH, with pH differences becoming apparent after approximately 5 min and a plateau reached after 30 min (**Figure 4A, B, Supporting Figure 4A, B**), This behavior is consistent with the solution and microparticle experiments and confirms that the encapsulated dye remained accessible to proton exchange after immobilization in the alginate network.

**Figure 4.**
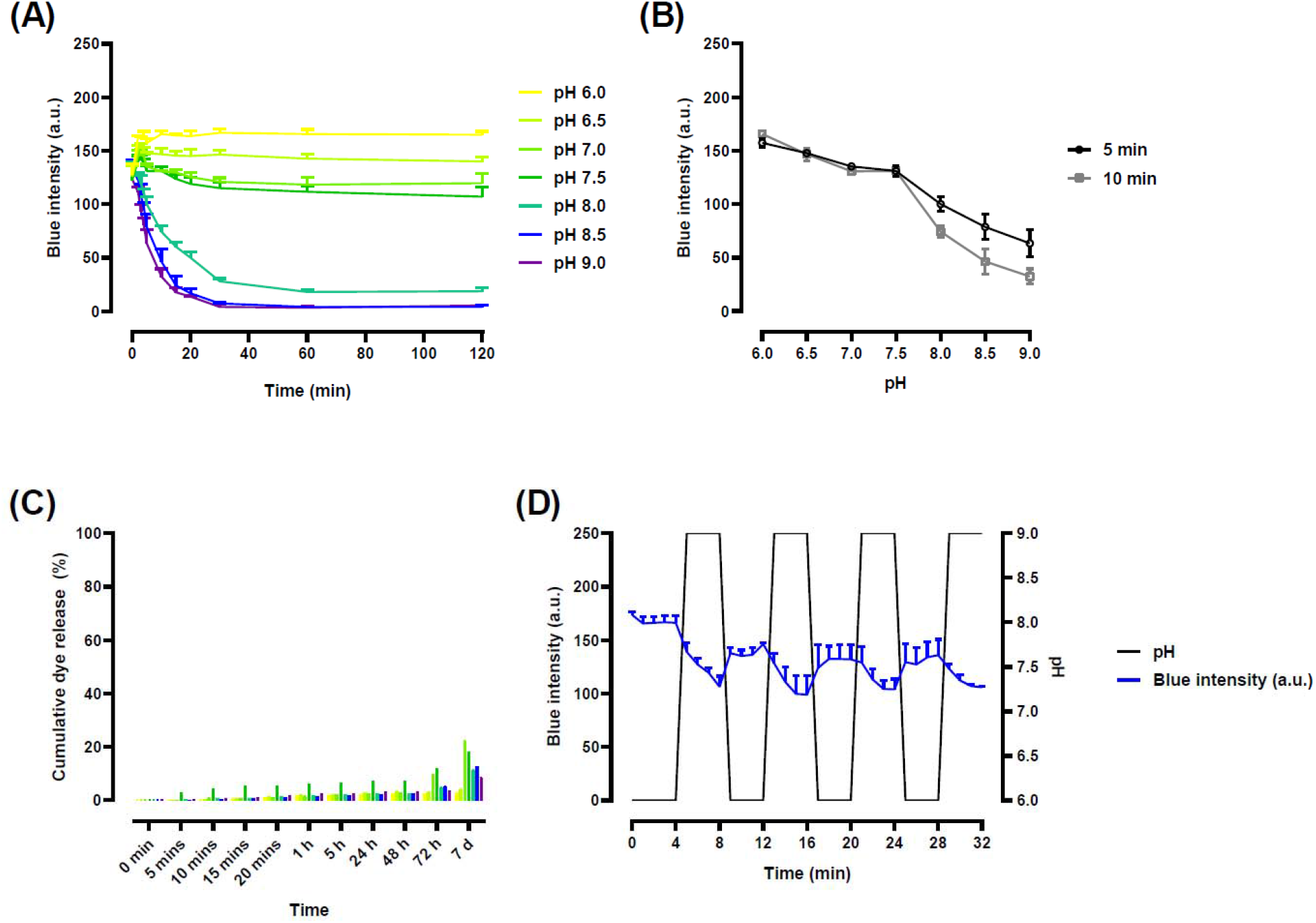
Colorimetric response of HPTS-loaded microparticles in calcium alginate hydrogels. **(A)** Kinetics of blue channel (B) intensity over 120 min following immersion in buffers at pH 6.0–9.0. **(B)** Blue channel intensity as a function of pH at 5 min and 10 min of buffer exposure, extracted from (A), showing pH-dependent separation of the signal at both incubation times. **(C)** Cumulative dye release (%) from HPTS-loaded hydrogels, quantified by fluorescence at 413 nm (HPTS isosbestic wavelength) in isotonic buffers at pH 6.0–9.0 (MES for pH 6.0–6.5; TBS for pH 7.0–9.0) at room temperature, monitored from 0 min to 7 days. **(D)** Reversibility of the blue channel intensity upon sequential exposure to pH 6.0 and pH 9.0 buffers over 32 min. Data are presented as mean ± SD (n = 3).

A similar pH-dependent decrease in blue intensity was observed *ex vivo* when the hydrogels were applied to full-thickness excisional dorsal mouse wounds (**Supporting Figure 4C, D**). The decrease in B value between the *in vitro* and *ex vivo* experiments indicates that background color from exposed wound tissue had limited impact on the hydrogel readout under the imaging conditions used here. This is an important translational point, because wound tissue presents a visually heterogeneous background that could otherwise interfere with RGB-based quantification. ^56^ The high density of HPTS-loaded microparticles within the dressing likely contributed to signal robustness by ensuring that the hydrogel dominated the measured color. Together, these findings show that the pH-responsive hydrogel remains analytically readable not only in controlled buffer systems but also in a more complex, wound-like environment.

The kinetics of the hydrogel signal was assessed by monitoring the B channel for up to 120 min after incubation in buffers. pH-dependent differences in blue intensity emerged after 5 min (**Figure 4B**), and the hydrogels developed progressively stronger yellow coloration at higher pH over time. As proton diffusion proceeded through the porous alginate matrix, the signal intensified and approached a plateau at approximately 30 min (**Figure 4A).** This behavior indicates that the response kinetics are governed both by the intrinsic acid–base equilibrium of HPTS and by the time required for buffer penetration and proton transport within the hydrogel network. Notably, the separation between pH conditions became apparent well before full equilibration, indicating that diagnostically useful information can be obtained after relatively short contact times. This is an advantage for wound applications, where prolonged measurement is often impractical. Over the 120 min window, the hydrogels maintained a stable and distinguishable colorimetric response at each pH, supporting the robustness of the readout during measurement (**Figure 4A**).

Reversibility was then examined by sequentially transferring the same hydrogels between pH 6.0 and pH 9.0 buffers, switching the buffer every 4 min while monitoring the response continuously (**Figure 4D**). Upon switching from pH 6.0 to pH 9.0, the B value decreased rapidly toward a lower plateau, whereas returning the hydrogels to pH 6.0 restored the signal toward its initial level. Repeated cycles produced the same alternating behavior, demonstrating that the sensing response is reversible and driven by pH-dependent proton exchange rather than irreversible dye conversion. This reversibility is important for chronic wound monitoring, where wound pH can evolve over time: in diabetic wounds, pH fluctuates in response to treatments and due to infection. ^11, 16, 17, 57^

To evaluate long-term dye retention in the microparticle-containing hydrogel, we monitored HPTS release into the surrounding buffer over 7 days by measuring fluorescence at 413 nm, the isosbestic wavelength of HPTS. Cumulative dye release remained low across all pH conditions through 72 h, increasing to 22% at pH 7.0 over 7 days (**Figure 4C**). Visible hydrogel degradation and clouding of solutions after 5 days points to potential hydrogel stability issues in solution over long incubation times as opposed to dye release. To prolong shelf-life, the hydrogel may require storage in calcium-containing media that increase the stability of calcium-crosslinked alginate hydrogels. ^58–63^

### 3.4. *In vivo* pH sensing in a murine wound model

For the *in vivo* proof-of-concept, we used a full-thickness dorsal wound model in mice. This model is widely established for wound healing studies. ^64, 65^ The geometry of the hydrogel was adapted for this model (**Supporting Figure 5A**). Relative to *the in vitro* and *ex vivo* hydrogels, the *in vivo* formulation used a higher microparticle loading and was cast as a thin film between microscope slides to produce flatter wound-sized patches. This change in geometry improved contact with the wound bed and enabled more reproducible imaging under controlled lighting after wound exposure. The hydrogel formulation adapted for use in full-thickness mouse wounds exhibited a preserved colorimetric pH-sensing profile (**Supporting Figure 5B, 5C**).

In healthy male C57BL/6 mice, the hydrogel was applied to the right wound at 0, 1, 3, and 24 h post-wounding, while wound fluid from the left wound was collected on filter paper at matching time points. In wound fluid collected from filter paper, diluted in saline, and analyzed with a commercial pH meter, we observed a slightly alkaline and increasing pH over the observed period **(Figure 5C**, **Supporting Table 1)**, consistent with the early hemostatic/inflammatory phase of acute full-thickness wound healing, during which increased microvascular permeability allows plasma and interstitial fluid (pH ≈ 7.4) to flood the wound bed. ^22^ This is distinct from the progressive acidification associated with the later proliferative phase, which occurs over days rather than hours. ^18, 19^ Similar pH values were observed in full-tickness mouse wounds in the literature. ^66–68^ Due to the significant pH difference in wound fluid at 0 and 24 h after wounding (**Figure 5A, B and Supporting Table 1**), we selected these time points to validate the pH-sensing hydrogel (**Figure 5C**). The hydrogel was placed on the wound for 10 min with a 180 degree rotation after 5 min, and its color was analyzed with the same smartphone setup as previously described. As the reference method indicated a wound fluid pH increase over 24 h (**Supporting Table 1**), we observed a significant decrease in B channel intensity over the same time period (**Figure 5C, Supporting Figure 6C)**. These reults demonstrate that the hydrogel was capable of detecting a difference in wound pH, and establish the *in vivo* proof-of-concept of using the colorimetric bandage for wound monitoring. The sensing chemistry remained unchanged relative to the *in vitro* and *ex vivo* formulations (only the geometry and microparticle loading was adapted to improve wound compatibility, **Supporting Figures 4A, 5A**), supporting the translational potential of the HPTS-microparticle-alginate system.

**Figure 5.**
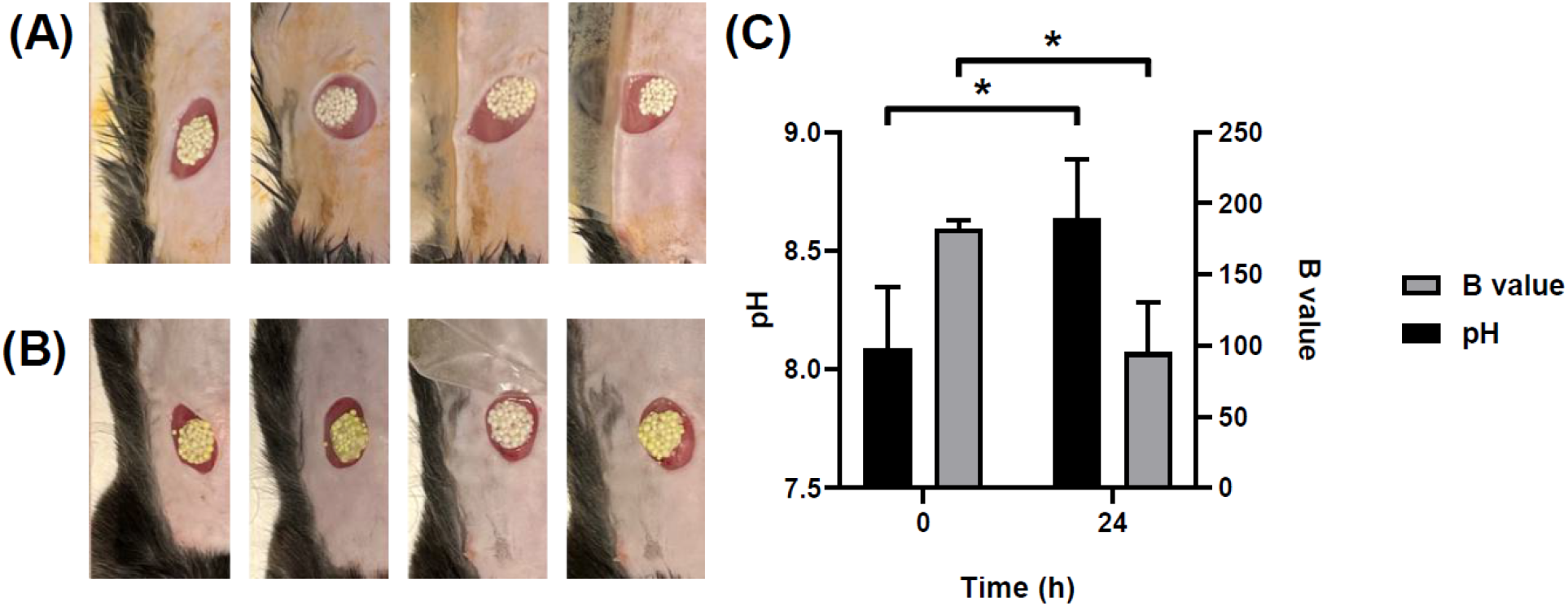
*In vivo* testing of pH-sensing colorimetric hydrogel. (A) Representative baseline (0 h) images of the hydrogel-covered wound right after wounding, (B) Representative images of the hydrogel-covered wound 24 h after wounding, (C) Wound fluid pH (left Y-axis, black bars; measured from filter paper collections) and corresponding hydrogel B values (right Y-axis, grey bars) at 0 h and 24 h (n = 4). All data are presented as mean ± SD; *p < 0.05.

## 4. Conclusion

In this study, we developed a colorimetric pH-sensing wound dressing that converts local wound pH into a quantitative optical signal readable with a conventional smartphone camera. By adsorbing the pH-sensitive dye HPTS onto cationic microparticles and encapsulating them within a calcium-crosslinked alginate hydrogel, we obtained a dressing whose blue-channel intensity decreased with increasing pH across the clinically relevant range of pH 6.0 to 9.0. The signal developed within minutes, remained stable over the measurement window, and was reversible across repeated acidic-to-alkaline transitions, indicating that the response reflects a dynamic acid–base equilibrium rather than irreversible dye conversion. This combination of speed, stability, and reversibility is well suited to longitudinal monitoring, where wound pH may shift as healing progresses or stalls.

Importantly, the colorimetric readout was preserved as the platform was developed from buffered dye solutions to microparticle suspensions, to encapsulated hydrogels. The pH-sensing hydrogel was successfully validated in a full-thickness mouse wound *in vivo*. The *in vivo* proof-of-concept demonstrated that rising wound pH was mirrored by a corresponding decrease in blue-channel intensity, and that the dense microparticle loading allows the dressing to quantify wound pH *in situ* despite the heterogeneous background of wound tissue. The adaptation of the hydrogel geometry for *in vivo* use in mice suggests that the HPTS-microparticle-alginate system can be reformatted for direct wound application in different geometries to address wound size differences in clinical practice. The combination of a pH-sensitive colorimetric hydrogel with a conventional smartphone detector offers accessible wound pH monitoring at the point-of-care.

## Acknowledgements

The authors thank Isabelle Caron, a technician at the In Vivo platform at IRIC, and Gabriela Manrique for their support with the *in vivo* studies. KC gratefully acknowledges support from the Bourse du centenaire of Université de Montréal.

---

**Table of Content figure.**
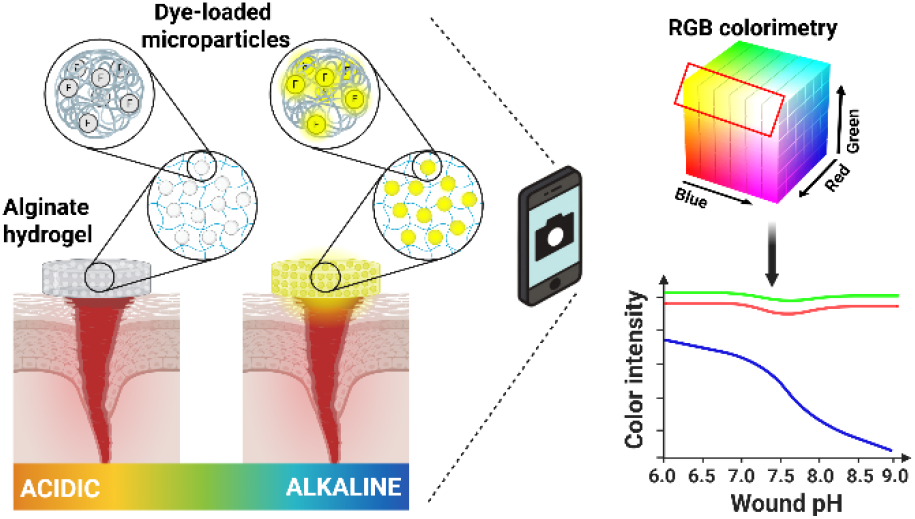
A colorimetric hydrogel dressing enables point-of-care wound pH monitoring via smartphone RGB analysis.

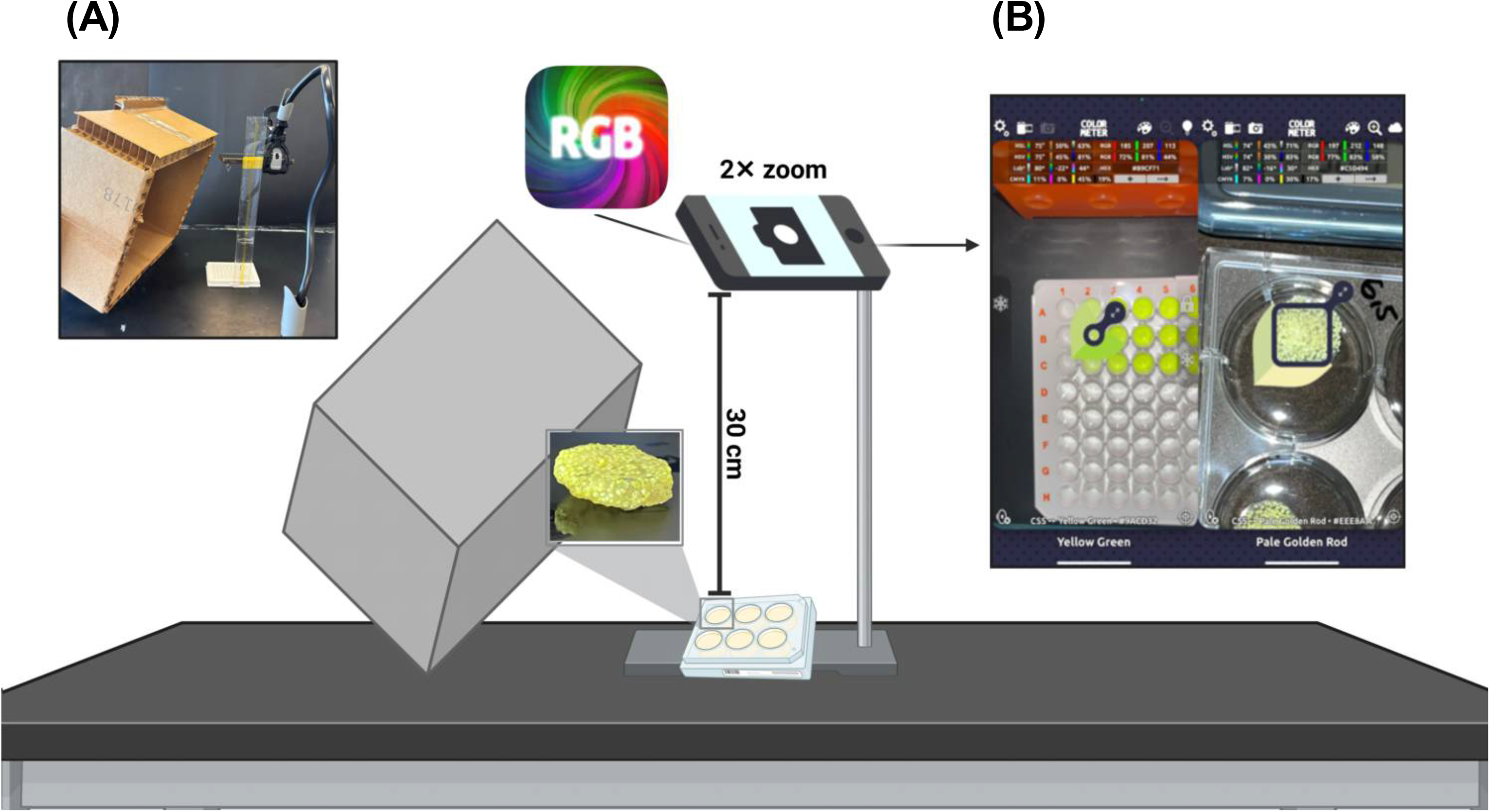

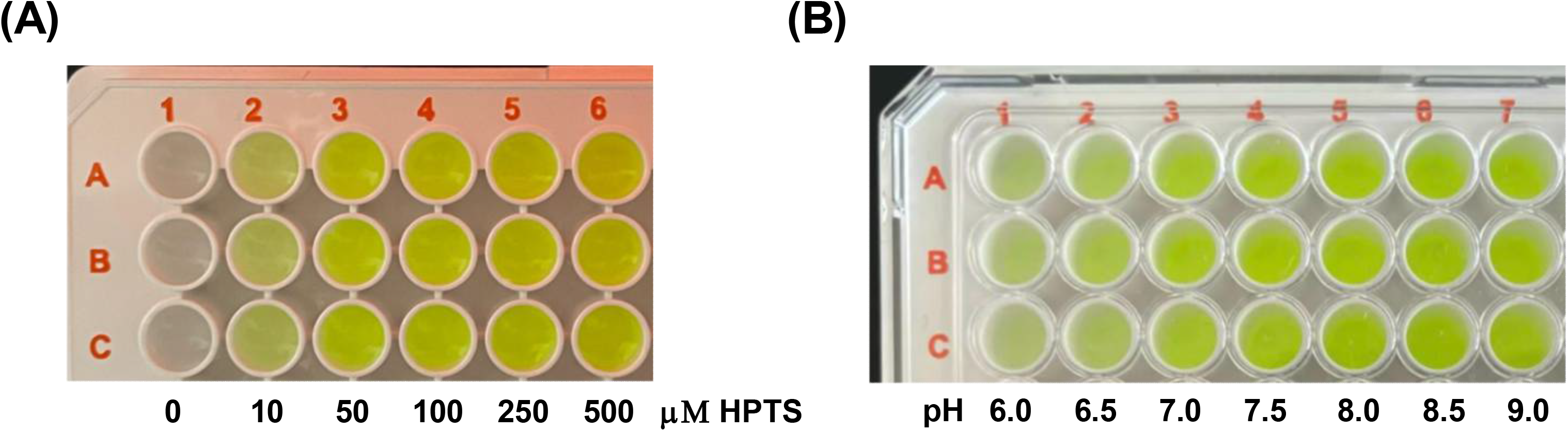

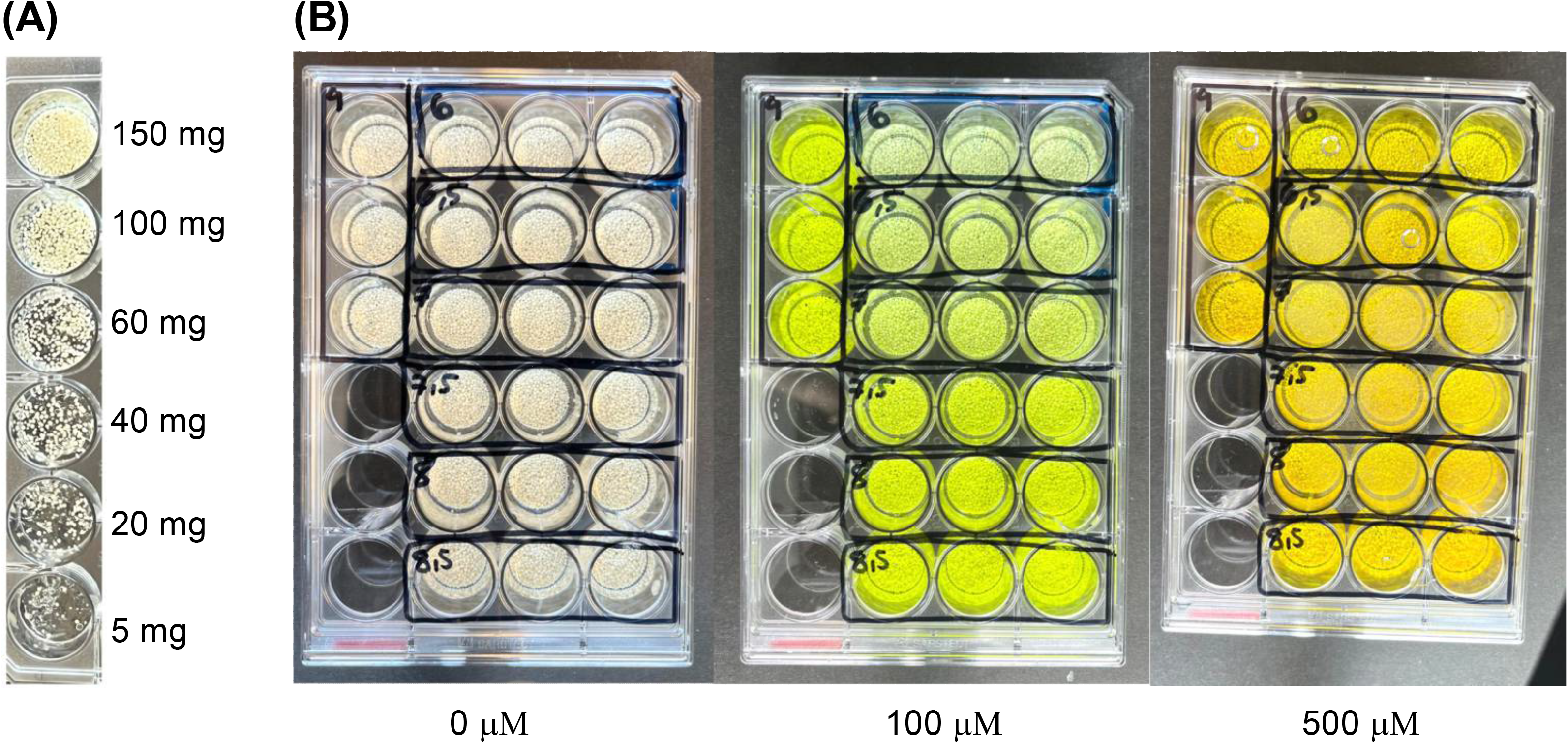

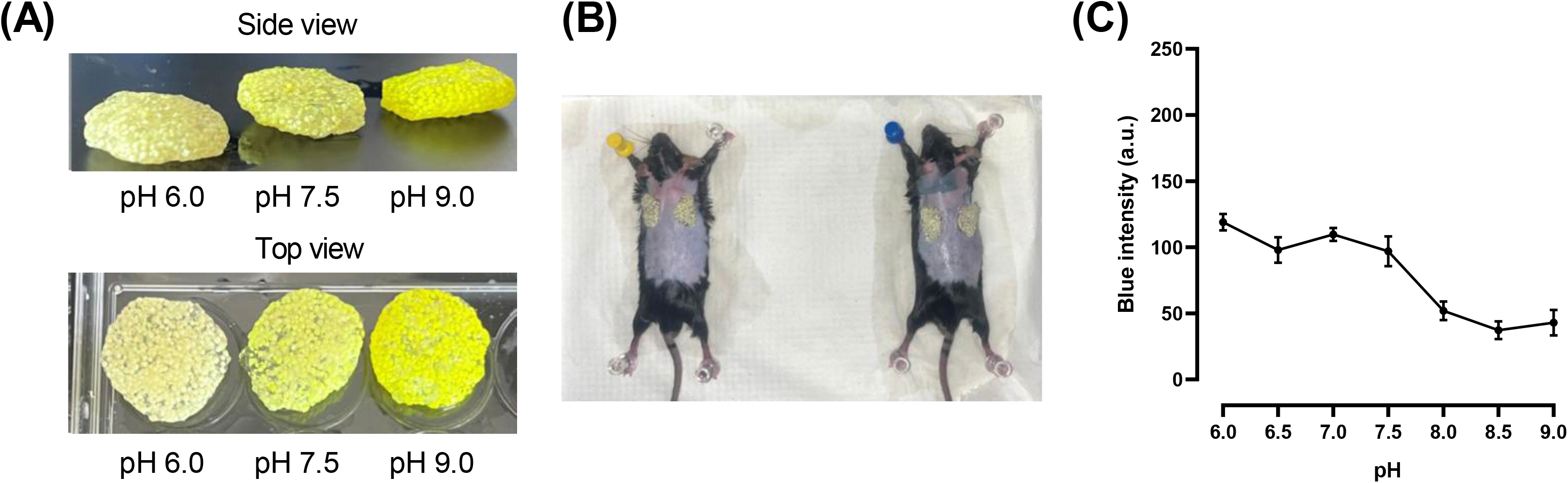

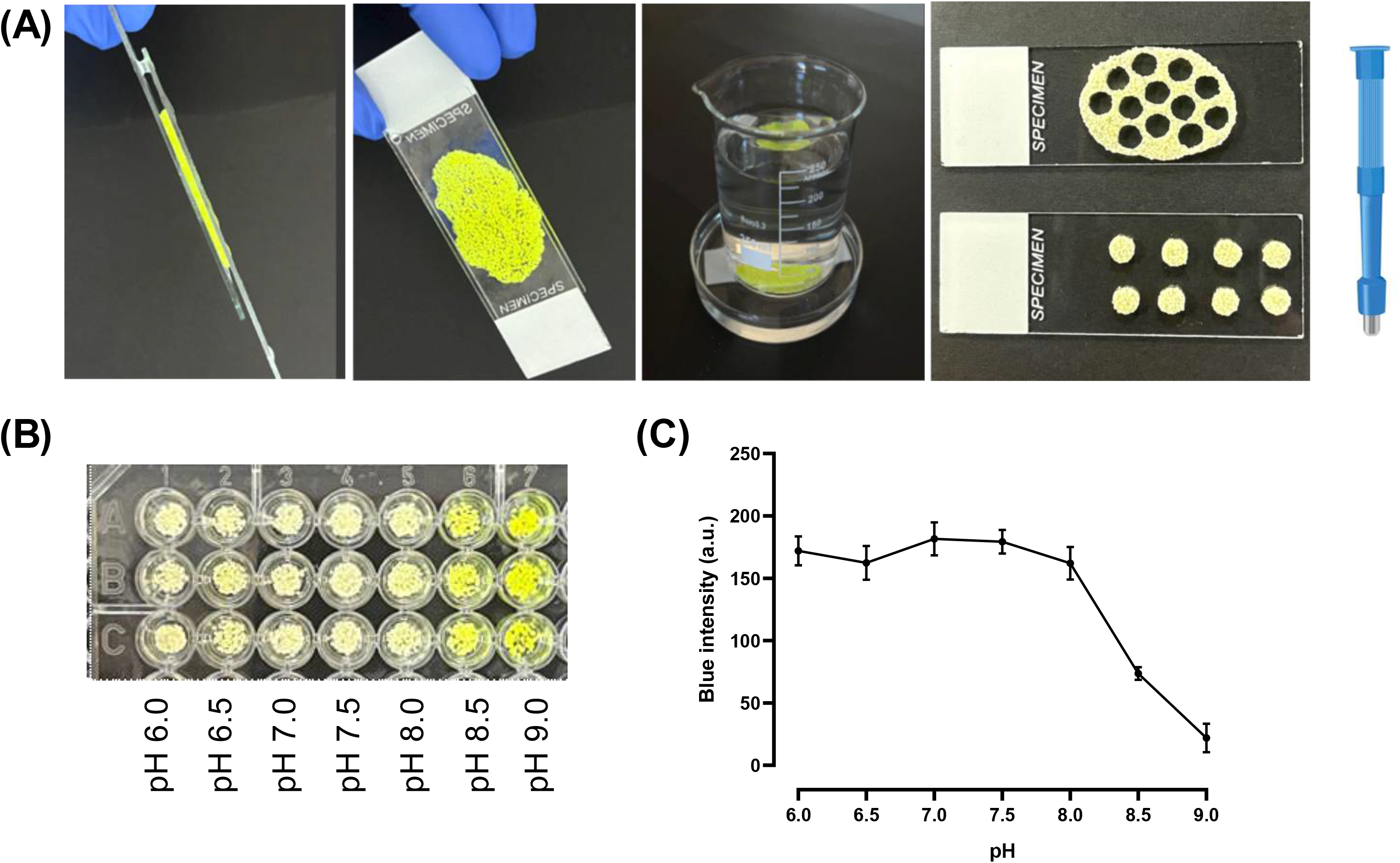

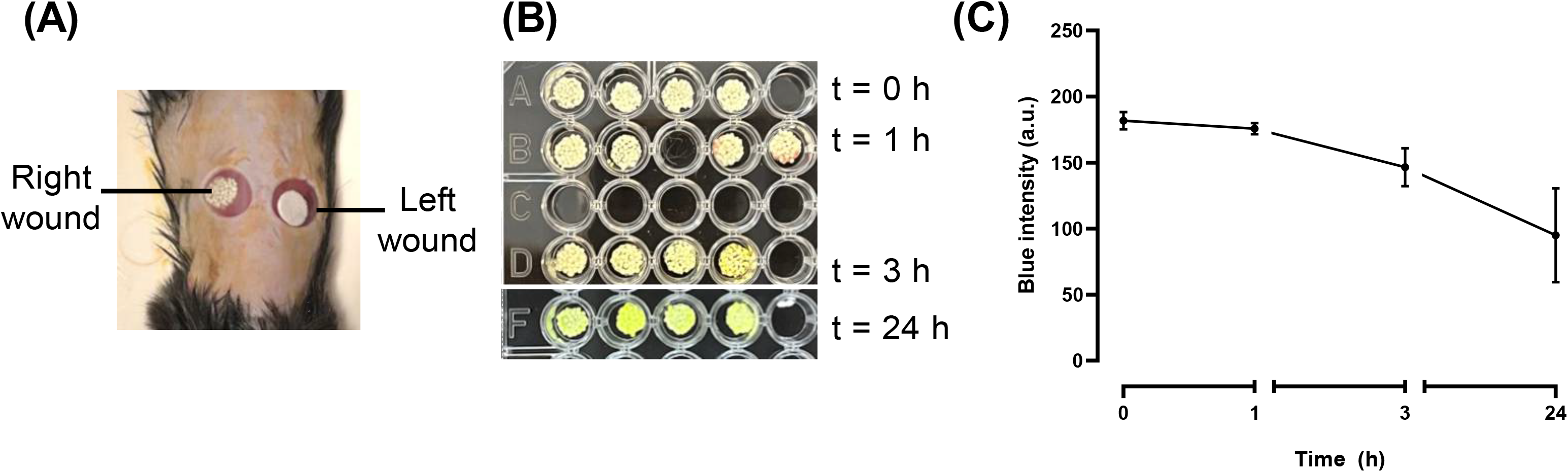

## References

(1) Armstrong, D. G.; Tan, T. W.; Boulton, A. J. M.; Bus, S. A. Diabetic Foot Ulcers: A Review. Jama 2023, 330 (1), 62–75. DOI: 10.1001/jama.2023.10578.

(2) Armstrong David, G.; Boulton Andrew, J. M.; Busr Sicco, A. Diabetic Foot Ulcers and Their Recurrence. New England Journal of Medicine 2017, 376 (24), 2367–2375. DOI: 10.1056/NEJMra1615439

(3) McDermott, K.; Fang, M.; Boulton, A. J. M.; Selvin, E.; Hicks, C. W. Etiology, Epidemiology, and Disparities in the Burden of Diabetic Foot Ulcers. Diabetes Care 2022, 46 (1), 209–221. DOI: 10.2337/dci22-0043.

(4) Oyibo, S. O.; Jude, E. B.; Tarawneh, I.; Nguyen, H. C.; Harkless, L. B.; Boulton, A. J. M. A Comparison of Two Diabetic Foot Ulcer Classification Systems: The Wagner and the University of Texas wound classification systems. Diabetes Care 2001, 24 (1), 84–88. DOI: 10.2337/diacare.24.1.84.

(5) Cerqueira, L. O.; Duarte, E. G.; Barros, A. L. S.; Cerqueira, J. R.; de Araújo, W. J. B. WIfI classification: the Society for Vascular Surgery lower extremity threatened limb classification system, a literature review. J Vasc Bras 2020, 19, e20190070. DOI: 10.1590/1677-5449.190070.

(6) Santema, T. B.; Lenselink, E. A.; Balm, R.; Ubbink, D. T. Comparing the Meggitt-Wagner and the University of Texas wound classification systems for diabetic foot ulcers: inter-observer analyses. Int Wound J 2016, 13 (6), 1137–1141. DOI: 10.1111/iwj.12429.

(7) Monteiro-Soares, M.; Hamilton, E. J.; Russell, D. A.; Srisawasdi, G.; Boyko, E. J.; Mills, J. L.; Jeffcoate, W.; Game, F. Guidelines on the classification of foot ulcers in people with diabetes (IWGDF 2023 update). Diabetes/Metabolism Research and Reviews 2024, 40 (3), e3648. DOI: 10.1002/dmrr.3648.

(8) Fu, T.; Stupnitskaia, P.; Matoori, S. Next-Generation Diagnostic Wound Dressings for Diabetic Wounds. ACS Measurement Science Au 2022, 2 (5), 377–384. DOI: 10.1021/acsmeasuresciau.2c00023.

(9) Qi, M.; Han, Y.; Zhang, W.; Liu, Y.; Jiang, D.; Wu, Z.; Xu, M.; Fu, J.; Li, B. Advances and future perspectives in hydrogel-based sensing technologies: a comprehensive review. Nanotechnology 2025, 36 (34). DOI: 10.1088/1361-6528/adf8f3.

(10) Tatum, O. L.; Dowd, S. E. Wound Healing Finally Enters the Age of Molecular Diagnostic Medicine. Adv Wound Care (New Rochelle) 2012, 1 (3), 115–119. DOI: 10.1089/wound.2011.0303.

(11) Wang, Y.; Miao, F.; Bai, J.; Wang, Z.; Qin, W. An observational study of the pH value during the healing process of diabetic foot ulcer. Journal of Tissue Viability 2024, 33 (2), 208–214. DOI: 10.1016/j.jtv.2024.03.015.

(12) McArdle, C.; Lagan, K. M.; McDowell, D. A. The pH of wound fluid in diabetic foot ulcers --the way forward in detecting clinical infection? Curr Diabetes Rev 2014, 10 (3), 177–181. DOI: 10.2174/1573399810666140609143217.

(13) Louisa, R. B.; Charne, N. M.; Richard, J. S.; Andrea, M. B. M.; Geoff, S.; William, M. The pH of wounds during healing and infection: a descriptive literature review. Wound Practice and Research 2017, 25 (2).

(14) Schreml, S.; Szeimies, R. M.; Karrer, S.; Heinlin, J.; Landthaler, M.; Babilas, P. The impact of the pH value on skin integrity and cutaneous wound healing. J Eur Acad Dermatol Venereol 2010, 24 (4), 373–378. DOI: 10.1111/j.1468-3083.2009.03413.x.

(15) Sim, P.; Strudwick, X. L.; Song, Y.; Cowin, A. J.; Garg, S. Influence of Acidic pH on Wound Healing In Vivo: A Novel Perspective for Wound Treatment. In International Journal of Molecular Sciences, 2022; Vol. 23.

(16) Tricou, L.-P.; Al-Hawat, M.-L.; Cherifi, K.; Manrique, G.; Freedman, B. R.; Matoori, S. Wound pH-Modulating Strategies for Diabetic Wound Healing. Advances in Wound Care 2024, 13 (9), 446–462. DOI: 10.1089/wound.2023.0129.

(17) Rembe, J.-D.; Witte, M.; Ertas, N.; Dissemond, J.; Garabet, W.; Hovhannisyan, M.; Beckamp, K.; Schelzig, H.; Wagenhäuser, M. U.; Stuermer, E. K. pH profiling reveals progressive wound acidification during healing and higher pH in chronic non-healing wounds: a prospective, multicenter cohort study. Scientific Reports 2026, 16 (1), 10522. DOI: 10.1038/s41598-026-45000-7.

(18) Schneider, L. A.; Korber, A.; Grabbe, S.; Dissemond, J. Influence of pH on wound-healing: a new perspective for wound-therapy? Arch Dermatol Res 2007, 298 (9), 413–420. DOI: 10.1007/s00403-006-0713-x.

(19) Percival, S. L.; McCarty, S.; Hunt, J. A.; Woods, E. J. The effects of pH on wound healing, biofilms, and antimicrobial efficacy. Wound Repair Regen 2014, 22 (2), 174–186. DOI: 10.1111/wrr.12125.

(20) Wu, H.; Yin, Y.; Hu, X.; Peng, C.; Liu, Y.; Li, Q.; Huang, W.; Huang, Q. Effects of Environmental pH on Macrophage Polarization and Osteoimmunomodulation. ACS Biomaterials Science & Engineering 2019, 5 (10), 5548–5557. DOI: 10.1021/acsbiomaterials.9b01181.

(21) Jones, E. M.; Cochrane, C. A.; Percival, S. L. The Effect of pH on the Extracellular Matrix and Biofilms. Adv Wound Care (New Rochelle) 2015, 4 (7), 431–439. DOI: 10.1089/wound.2014.0538.

(22) Kruse, C. R.; Singh, M.; Targosinski, S.; Sinha, I.; Sørensen, J. A.; Eriksson, E.; Nuutila, K. The effect of pH on cell viability, cell migration, cell proliferation, wound closure, and wound reepithelialization: In vitro and in vivo study. Wound Repair and Regeneration 2017, 25 (2), 260–269. DOI: 10.1111/wrr.12526.

(23) Sharpe, J. R.; Harris, K. L.; Jubin, K.; Bainbridge, N. J.; Jordan, N. R. The effect of pH in modulating skin cell behaviour. British Journal of Dermatology 2009, 161 (3), 671–673. DOI: 10.1111/j.1365-2133.2009.09168.x.

(24) Schreml, S.; Meier, R. J.; Kirschbaum, M.; Kong, S. C.; Gehmert, S.; Felthaus, O.; Küchler, S.; Sharpe, J. R.; Wöltje, K.; Weiß, K. T.;, et al. Luminescent dual sensors reveal extracellular pH-gradients and hypoxia on chronic wounds that disrupt epidermal repair. Theranostics 2014, 4 (7), 721–735. DOI: 10.7150/thno.9052.

(25) Al-Hawat, M. L.; Cherifi, K.; Tricou, L. P.; Lamontagne, S.; Tran, M.; Ngu, A. C. Y.; Manrique, G.; Guirguis, N.; Machuca-Parra, A. I.; Matoori, S. Fluorescent pH-sensing bandage for point-of-care wound diagnostics. Aggregate 2024, 5 (2). DOI: 10.1002/agt2.472.

(26) Yang, J.; Wang, K.; Xu, H.; Yan, W.; Jin, Q.; Cui, D. Detection platforms for point-of-care testing based on colorimetric, luminescent and magnetic assays: A review. Talanta 2019, 202, 96–110. DOI: 10.1016/j.talanta.2019.04.054.

(27) Berg, B.; Cortazar, B.; Tseng, D.; Ozkan, H.; Feng, S.; Wei, Q.; Chan, R. Y.-L.; Burbano, J.; Farooqui, Q.; Lewinski, M.;, et al. Cellphone-Based Hand-Held Microplate Reader for Point-of-Care Testing of Enzyme-Linked Immunosorbent Assays. ACS Nano 2015, 9 (8), 7857–7866. DOI: 10.1021/acsnano.5b03203.

(28) Cherifi, K.; Toupchinejad, F.; Christodoulopoulos, K.; Kizilkaya, A.; Matoori, S. Colorimetric detection methods of pH-sensing wound dressing for point-of-care wound diagnostics. BioRxiv, 2024.

(29) Gu, H.; Sun, X.; Bao, H.; Feng, X.; Chen, Y. Optically pH-Sensing in smart wound dressings towards real-time monitoring of wound states: A review. Analytica Chimica Acta 2025, 1350, 343808. DOI: 10.1016/j.aca.2025.343808.

(30) Liu, L.; Li, X.; Nagao, M.; Elias, A. L.; Narain, R.; Chung, H.-J. A pH-Indicating Colorimetric Tough Hydrogel Patch towards Applications in a Substrate for Smart Wound Dressings. In Polymers, 2017; Vol. 9.

(31) Zheng, X. T.; Zhong, Y.; Chu, H. E.; Yu, Y.; Zhang, Y.; Chin, J. S.; Becker, D. L.; Su, X.; Loh, X. J. Carbon Dot-Doped Hydrogel Sensor Array for Multiplexed Colorimetric Detection of Wound Healing. ACS Applied Materials & Interfaces 2023, 15 (14), 17675–17687. DOI: 10.1021/acsami.3c01185.

(32) Soda, Y.; Bakker, E. Quantification of Colorimetric Data for Paper-Based Analytical Devices. ACS Sensors 2019, 4 (12), 3093–3101. DOI: 10.1021/acssensors.9b01802.

(33) Chandra, A.; Prasad, S.; Iuele, H.; Colella, F.; Rizzo, R.; D’Amone, E.; Gigli, G.; Del Mercato, L. L. Highly Sensitive Fluorescent pH Microsensors Based on the Ratiometric Dye Pyranine Immobilized on Silica Microparticles. Chemistry 2021, 27 (53), 13318–13324. DOI: 10.1002/chem.202101568.

(34) Clement, N. R.; Gould, J. M. Pyranine (8-hydroxy-1,3,6-pyrenetrisulfonate) as a probe of internal aqueous hydrogen ion concentration in phospholipid vesicles. Biochemistry 1981, 20 (6), 1534–1538. DOI: 10.1021/bi00509a019.

(35) Kano, K.; Fendler, J. H. Pyranine as a sensitive pH probe for liposome interiors and surfaces. pH gradients across phospholipid vesicles. Biochim Biophys Acta 1978, 509 (2), 289–299. DOI: 10.1016/0005-2736(78)90048-2.

(36) Nandi, R.; Amdursky, N. The Dual Use of the Pyranine (HPTS) Fluorescent Probe: A Ground-State pH Indicator and an Excited-State Proton Transfer Probe. Accounts of Chemical Research 2022, 55 (18), 2728–2739. DOI: 10.1021/acs.accounts.2c00458.

(37) Panzarasa, G.; Osypova, A.; Toncelli, C.; Buhmann, M. T.; Rottmar, M.; Ren, Q.; Maniura-Weber, K.; Rossi, R. M.; Boesel, L. F. The pyranine-benzalkonium ion pair: A promising fluorescent system for the ratiometric detection of wound pH. Sensors and Actuators B: Chemical 2017, 249, 156–160. DOI: 10.1016/j.snb.2017.04.045.

(38) Thomas, J. V.; Brimijoin, M. R.; Neault, T. R.; Brubaker, R. F. The fluorescent indicator pyranine is suitable for measuring stromal and cameral pH in vivo. Exp Eye Res 1990, 50 (3), 241–249. DOI: 10.1016/0014-4835(90)90208-c.

(39) GmbH, W. M. ColorMeter RGB Colorimeter: Color Picker & Measurement. 2013–2022. https://apps.apple.com/us/app/colormeter-rgb-colorimeter/id713258885 (accessed).

(40) Huang, Y.-Y.; Lin, C.-W.; Cheng, N.-C.; Cazzell, S. M.; Chen, H.-H.; Huang, K.-F.; Tung, K.-Y.; Huang, H.-L.; Lin, P.-Y.; Perng, C.-K.;, et al. Effect of a Novel Macrophage-Regulating Drug on Wound Healing in Patients With Diabetic Foot Ulcers: A Randomized Clinical Trial. JAMA Network Open 2021, 4 (9), e2122607–e2122607. DOI: 10.1001/jamanetworkopen.2021.22607.

(41) Charbonier, F.; Indana, D.; Chaudhuri, O. Tuning Viscoelasticity in Alginate Hydrogels for 3D Cell Culture Studies. Current Protocols 2021, 1 (5), e124. DOI: 10.1002/cpz1.124.

(42) Smiell, J. M.; Wieman, T. J.; Steed, D. L.; Perry, B. H.; Sampson, A. R.; Schwab, B. H. Efficacy and safety of becaplermin (recombinant human platelet-derived growth factor-BB) in patients with nonhealing, lower extremity diabetic ulcers: a combined analysis of four randomized studies. Wound Repair Regen 1999, 7 (5), 335–346. DOI: 10.1046/j.1524-475x.1999.00335.x.

(43) Ramsay, S.; Cowan, L.; Davidson, J. M.; Nanney, L.; Schultz, G. Wound samples: moving towards a standardised method of collection and analysis. International Wound Journal 2016, 13 (5), 880–891. DOI: 10.1111/iwj.12399.

(44) Zheng, X. T.; Yang, Z.; Sutarlie, L.; Thangaveloo, M.; Yu, Y.; Salleh, N. A. B. M.; Chin, J. S.; Xiong, Z.; Becker, D. L.; Loh, X. J.;, et al. Battery-free and AI-enabled multiplexed sensor patches for wound monitoring. Science Advances 9 (24), eadg6670. DOI: 10.1126/sciadv.adg6670.

(45) Matoori, S.; Bao, Y.; Schmidt, A.; Fischer, E. J.; Ochoa-Sanchez, R.; Tremblay, M.; Oliveira, M. M.; Rose, C. F.; Leroux, J. C. An Investigation of PS-b-PEO Polymersomes for the Oral Treatment and Diagnosis of Hyperammonemia. Small 2019, 15 (50), e1902347. DOI: 10.1002/smll.201902347.

(46) Yang, X.; Min, X.; Nan, X.; Gong, Z.; Chen, J.; Huang, W.; Sun, Y.; Zhou, M.; Wang, X.; Yi, X. A smartphone-assisted colorimetric nanoprobe for rapid detection of azodicarbonamide. Microchemical Journal 2025, 218, 115699. DOI: 10.1016/j.microc.2025.115699.

(47) Jacinto, C.; Maza Mejía, I.; Khan, S.; López, R.; Sotomayor, M. D. P. T.; Picasso, G. Using a Smartphone-Based Colorimetric Device with Molecularly Imprinted Polymer for the Quantification of Tartrazine in Soda Drinks. In Biosensors, 2023; Vol. 13, p 639.

(48) Steinegger, A.; Wolfbeis, O. S.; Borisov, S. M. Optical Sensing and Imaging of pH Values: Spectroscopies, Materials, and Applications. Chemical Reviews 2020, 120 (22), 12357–12489. DOI: 10.1021/acs.chemrev.0c00451.

(49) Abka-Khajouei, R.; Tounsi, L.; Shahabi, N.; Patel, A. K.; Abdelkafi, S.; Michaud, P. Structures, Properties and Applications of Alginates. Mar Drugs 2022, 20 (6). DOI: 10.3390/md20060364.

(50) Augst, A. D.; Kong, H. J.; Mooney, D. J. Alginate Hydrogels as Biomaterials. Macromolecular Bioscience 2006, 6 (8), 623–633. DOI: 10.1002/mabi.200600069.

(51) Gheorghita Puscaselu, R.; Lobiuc, A.; Dimian, M.; Covasa, M. Alginate: From Food Industry to Biomedical Applications and Management of Metabolic Disorders. Polymers (Basel) 2020, 12 (10). DOI: 10.3390/polym12102417.

(52) Matoori, S.; Veves, A.; Mooney, D. J. Advanced bandages for diabetic wound healing. Science Translational Medicine 2021, 13 (585), eabe4839. DOI: 10.1126/scitranslmed.abe4839.

(53) Freedman, B. R.; Hwang, C.; Talbot, S.; Hibler, B.; Matoori, S.; Mooney, D. J. Breakthrough treatments for accelerated wound healing. Sci Adv 2023, 9 (20). DOI: 10.1126/sciadv.ade7007.

(54) Freedman, B. R.; Mooney, D. J. Biomaterials to Mimic and Heal Connective Tissues. Advanced Materials 2019, 31 (19), 1806695. DOI: 10.1002/adma.201806695

(55) Freedman, B. R.; Uzun, O.; Luna, N. M. M.; Rock, A.; Clifford, C.; Stoler, E.; Östlund-Sholars, G.; Johnson, C.; Mooney, D. J. Degradable and Removable Tough Adhesive Hydrogels. Adv Mater 2021, 33 (17), e2008553. DOI: 10.1002/adma.202008553.

(56) Veredas, F. J.; Mesa, H.; Morente, L. Efficient detection of wound-bed and peripheral skin with statistical colour models. Medical & Biological Engineering & Computing 2015, 53 (4), 345–359. DOI: 10.1007/s11517-014-1240-0.

(57) Gethin, G.; O’Connor, G. M.; Abedin, J.; Newell, J.; Flynn, L.; Watterson, D.; O’Loughlin, A. Monitoring of pH and temperature of neuropathic diabetic and nondiabetic foot ulcers for 12 weeks: An observational study. Wound Repair and Regeneration 2018, 26 (2), 251–256. DOI: 10.1111/wrr.12628.

(58) Borgogna, M.; Skjåk-Bræk, G.; Paoletti, S.; Donati, I. On the initial binding of alginate by calcium ions. The tilted egg-box hypothesis. J Phys Chem B 2013, 117 (24), 7277–7282. DOI: 10.1021/jp4030766.

(59) Kuo, C. K.; Ma, P. X. Maintaining dimensions and mechanical properties of ionically crosslinked alginate hydrogel scaffolds in vitro. Journal of Biomedical Materials Research Part A 2008, 84A (4), 899–907. DOI: 10.1002/jbm.a.31375 (accessed 2026/07/31).

(60) Cao, L.; Lu, W.; Mata, A.; Nishinari, K.; Fang, Y. Egg-box model-based gelation of alginate and pectin: A review. Carbohydrate Polymers 2020, 242, 116389. DOI: 10.1016/j.carbpol.2020.116389.

(61) Grant, G. T.; Morris, E. R.; Rees, D. A.; Smith, P. J. C.; Thom, D. Biological interactions between polysaccharides and divalent cations: The egg-box model. FEBS Letters 1973, 32 (1), 195–198. DOI: 10.1016/0014-5793(73)80770-7.

(62) Makarova, A. O.; Derkach, S. R.; Khair, T.; Kazantseva, M. A.; Zuev, Y. F.; Zueva, O. S. Ion-Induced Polysaccharide Gelation: Peculiarities of Alginate Egg-Box Association with Different Divalent Cations. Polymers (Basel) 2023, 15 (5). DOI: 10.3390/polym15051243.

(63) Plazinski, W. Molecular basis of calcium binding by polyguluronate chains. Revising the egg-box model. J Comput Chem 2011, 32 (14), 2988–2995. DOI: 10.1002/jcc.21880.

(64) Castleberry, S. A.; Almquist, B. D.; Li, W.; Reis, T.; Chow, J.; Mayner, S.; Hammond, P. T. Self-Assembled Wound Dressings Silence MMP-9 and Improve Diabetic Wound Healing In Vivo. Adv Mater 2016, 28 (9), 1809–1817. DOI: 10.1002/adma.201503565.

(65) Nguyen, T. T.; Ding, D.; Wolter, W. R.; Pérez, R. L.; Champion, M. M.; Mahasenan, K. V.; Hesek, D.; Lee, M.; Schroeder, V. A.; Jones, J. I.;, et al. Validation of Matrix Metalloproteinase-9 (MMP-9) as a Novel Target for Treatment of Diabetic Foot Ulcers in Humans and Discovery of a Potent and Selective Small-Molecule MMP-9 Inhibitor That Accelerates Healing. Journal of Medicinal Chemistry 2018, 61 (19), 8825–8837. DOI: 10.1021/acs.jmedchem.8b01005.

(66) Mai, H.; Wang, Y.; Li, S.; Jia, R.; Li, S.; Peng, Q.; Xie, Y.; Hu, X.; Wu, S. A pH-sensitive near-infrared fluorescent probe with alkaline pK(a) for chronic wound monitoring in diabetic mice. Chem Commun (Camb) 2019, 55 (51), 7374–7377. DOI: 10.1039/c9cc02289a.

(67) Mirani, B.; Hadisi, Z.; Pagan, E.; Dabiri, S. M. H.; van Rijt, A.; Almutairi, L.; Noshadi, I.; Armstrong, D. G.; Akbari, M. Smart Dual-Sensor Wound Dressing for Monitoring Cutaneous Wounds. Advanced Healthcare Materials 2023, 12 (18), 2203233. DOI: 10.1002/adhm.202203233.

(68) Sharpe, J. R.; Booth, S.; Jubin, K.; Jordan, N. R.; Lawrence-Watt, D. J.; Dheansa, B. S. Progression of wound pH during the course of healing in burns. J Burn Care Res 2013, 34 (3), e201–208. DOI: 10.1097/BCR.0b013e31825d5569.

