## Supporting Information for "Colorimetric Hydrogel Dressing with Smartphone Detector for Point-of-Care Wound pH Monitoring"

Simon Matoori

Université de Montréal

2940 Chemin de Polytechnique, Montreal, QC H3T 1J4

**Supporting Information**

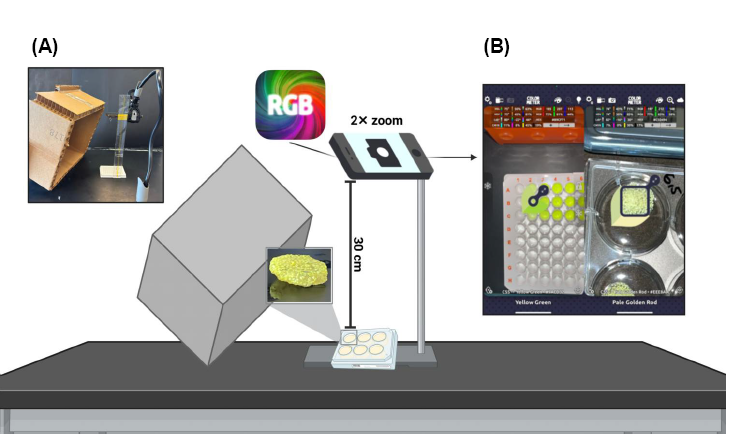

**Supporting Figure 1.** Smartphone-based RGB imaging setup. **(A)** Photo and schematics of the imaging setup used to quantify the colorimetric response of HPTS in solution, on microparticles, and in microparticle-loaded hydrogels. Samples were positioned on a rigid holder 30 cm below an iPhone 13 rear camera fixed at 2× zoom, with a black cardboard box held over the setup to minimize specular reflections from ambient lighting. **(B)** Representative ColorMeter App image of HPTS solutions in a 96-well plate, illustrating regions of interest (ROI) used to extract mean R, G, and B channel intensities.

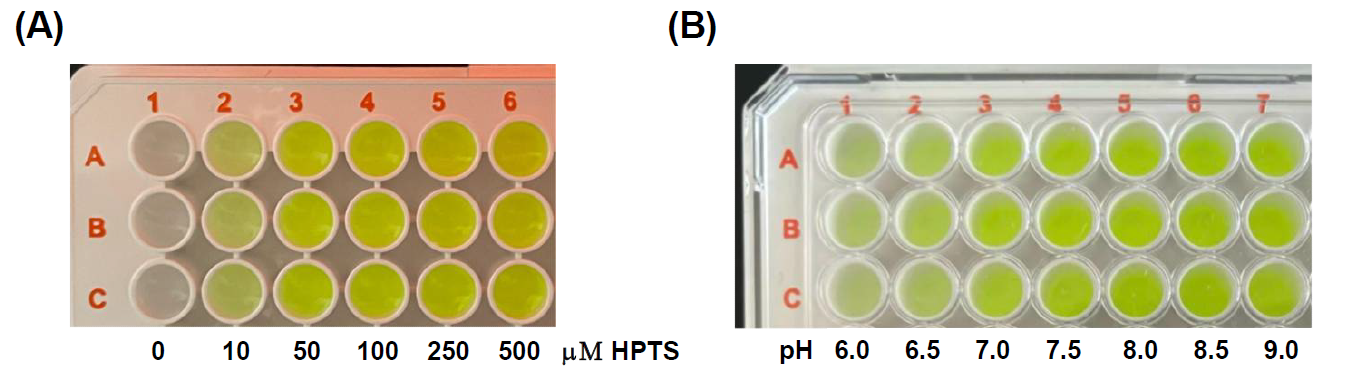

**Supporting Figure 2**. Colorimetric response of HPTS solutions to concentration and pH. **(A)** Representative photographs of HPTS solutions (0–500 µM) in PBS buffer (pH 7.4), showing a concentration-dependent increase in color intensity**. (B)** Representative photographs of HPTS solutions (100 µM) across pH 6.0–9.0, showing a pH-dependent colorimetric shift. Each row (Series 1–3) represents an independent replicate (n = 3). Images shown here were taken under ambient lighting for illustrative purposes only and do not reflect the controlled imaging conditions (Supporting Figure 1) used for quantitative RGB analysis.

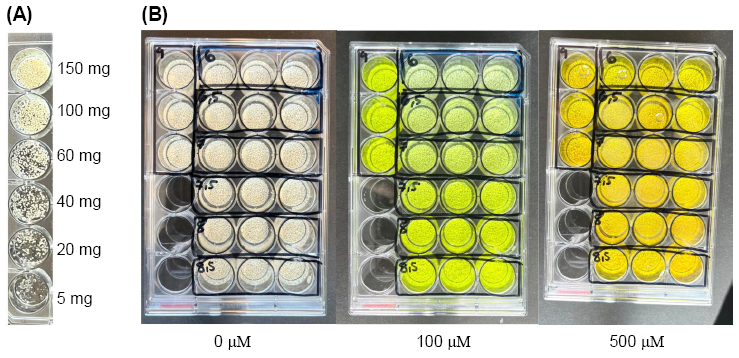

**Supporting Figure 3.** BTA microparticle loading and colorimetric response to HPTS. **(A)** Representative photographs of BTA microparticles at increasing mass (5–150 mg), used to select the 150 mg loading. **(B)** Representative photographs of BTA microparticles incubated with 0, 100, or 500 µM HPTS; the left column shows pH 9.0, and the right columns show pH 6.0–8.5 in 0.5 increments (top to bottom). Images were taken under ambient lighting for illustration only and do not reflect the controlled conditions (Supporting Figure 1) used for quantitative analysis.

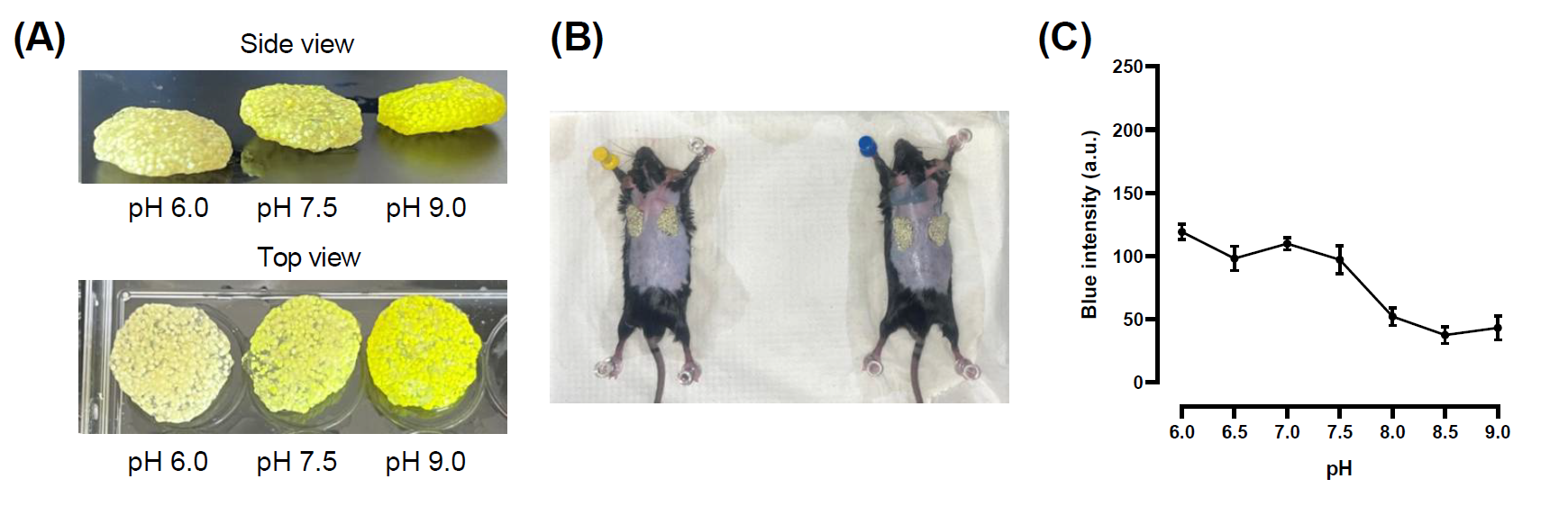

**Supporting Figure 4.** *Ex vivo* testing of the pH-sensing colorimetric hydrogel. **(A)** Representative photograph of the HPTS-loaded colorimetric hydrogel used for *ex vivo* testing. **(B)** Representative *ex vivo* wound setup, shown for 2 of 3 mice tested at pH 6.0. **(C)** Blue channel (B) intensity of hydrogels pre-equilibrated in buffers at pH 6.0–9.0 for 5 min and then placed onto excised full-thickness mouse wounds prior to colorimetry, showing a pH-dependent decrease in signal consistent with the *in vitro* response. Data are presented as mean ± SD (n = 3).

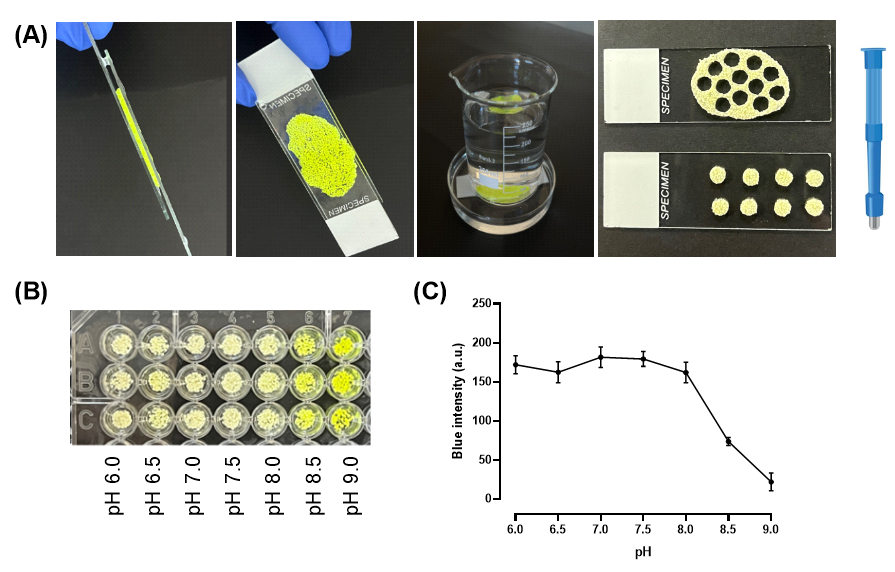

**Supporting Figure 5.** Preparation and testing of the microparticle-loaded hydrogels adapted to the mouse wound geometry. **(A)** Preparation of the hydrogels, including film casting between glass slides and biopsy punching into wound-sized patches**. (B)** Representative photographs of hydrogels equilibrated at pH 6.0–9.0, used to generate the standard curve. **(C)** Standard curve showing blue channel (B) intensity of the hydrogels as a function of pH, after 10 min incubation in MES buffer (pH 6.0–6.5) or TBS buffer (pH 7.0–9.0). Data are presented as mean ± SD (n = 3).

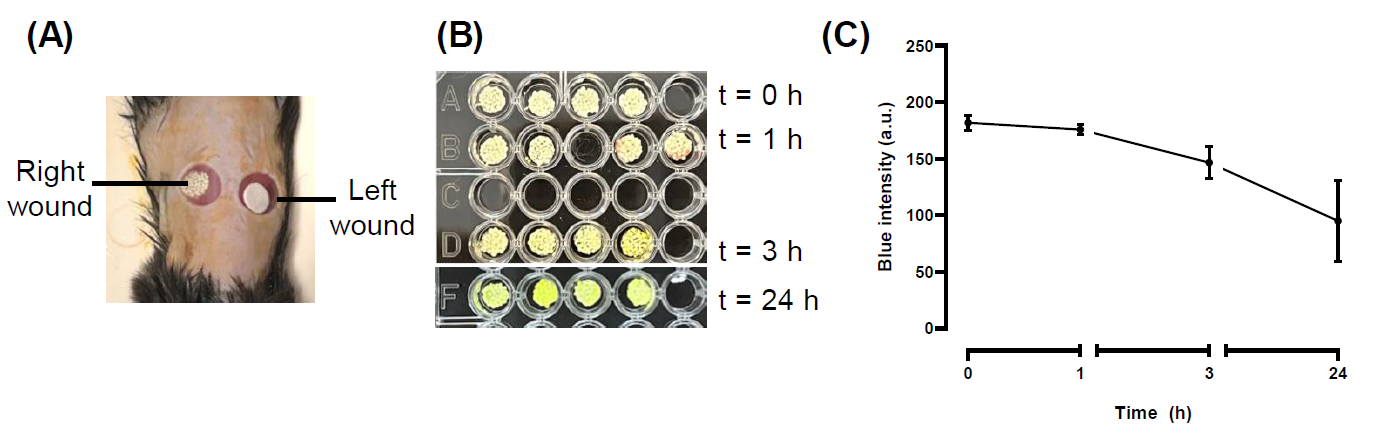

**Supporting Figure 6.** *In vivo* colorimetric response of the microparticle-loaded hydrogels. **(A)** Representative baseline images (0 h) showing the right wound covered with the pH‑sensing hydrogel and the left wound covered with the Whatman filter paper for wound fluid collection, **(B)** Photographs of the hydrogels recovered from mouse wounds at 0, 1, 3, and 24 h post-wounding. **(C)** Blue channel (B) intensity of the wound-applied hydrogels over time (0, 1, 3, and 24 h). Data are presented as mean ± SD (n = 4).

| Time after wounding (h) | Wound fluid volume sampled (µL) | Measured pH |
| --- | --- | --- |
| 0 | 2.88 ± 0.36 | 8.09 ± 0.26 |
| 1 | 2.78 ± 1.00 | 8.21 ± 0.11 |
| 3 | 3.00 ± 0.54 | 8.43 ± 0.09 |
| 24 | 5.33 ± 3.62 | 8.69 ± 0.19 |

**Supporting Table 1**. Wound fluid volume and pH measured at each time point in the murine wound model. Wound fluid volume and corresponding pH were measured at 0, 1, 3, and 24 h post-wounding using filter paper collection from the contralateral wound. Data are presented as mean ± SD (n = 4).
